# Cyclic-di-GMP induces STING-dependent ILC2 to ILC1 shift during innate type 2 lung inflammation

**DOI:** 10.1101/2020.10.19.345850

**Authors:** Kellen J. Cavagnero, Jana H. Badrani, Luay H. Naji, Michael B. Amadeo, Anthea S. Leng, Lee Diego Lacasa, Allyssa N. Strohm, Taylor A. Doherty

## Abstract

Type 2 inflammation is found in most forms of asthma, which may co-exist with recurrent viral infections, bacterial colonization, and host cell death. These processes drive the accumulation of intracellular cyclic-di-nucleotides such as cyclic-di-GMP (CDG). Group 2 innate lymphoid cells (ILC2s) are critical drivers of type 2 lung inflammation during fungal allergen exposure in mice; however, it is unclear how CDG regulates lung ILC responses during lung inflammation. Here, we show that CDG induced early airway type 1 interferon (IFN) production and dramatically suppressed both lung proliferating CD127+ST2+ ILC2s and *Alternaria*- and IL-33-induced lung inflammation. Further, transcriptomic analysis of CD127−ST2− Thy1.2+ ILCs, which were expanded and activated by CDG, revealed an ILC1 signature. CDG administration led to accumulation of IFNγ+ and T-bet+ ILC1s, as well as neutrophilia, independent of IL-18R, IL-12, and STAT6 but dependent on stimulator of interferon genes (STING) and partially dependent on type 1 IFN signaling. Thus, CDG potently suppresses ILC2-driven lung inflammation and promotes ILC1 responses. These results suggest potential therapeutic modulation of STING to suppress type 2 inflammation and/or increase anti-viral responses during respiratory infections.

## INTRODUCTION

Group 2 innate lymphoid cells (ILC2s) were identified in 2010 and have since been recognized for their essential role in orchestrating innate type 2 immune responses in murine asthma models (1-3). Airway exposure to the clinically-relevant fungal allergen *Alternaria alternata* (*Alt*) promotes rapid epithelial cell IL-33 secretion and ILC2 production of IL-5 and IL-13, which drive peribronchial eosinophilic inflammation (4). ILC2s belong to a greater family of ILCs that includes ILC1s, which are involved in protective immune responses against intracellular pathogens. While ILC2s express the key transcription factor GATA-3 and produce type 2 cytokines, ILC1s express T-bet and produce interferon-γ (IFNγ).

Insights into asthma pathogenesis have revealed associations with bacterial colonization (5), viral infections (6), as well as inappropriate cell death and mitochondrial stress (7). These processes drive accumulation of cyclic-di-nucleotide danger signals inside of host cells (8). Free intracellular viral and host DNA are converted to cyclic-di-nucleotides by the host cytosolic enzyme cyclic-GMP-AMP-synthase (9). One particular cyclic-di-nucleotide, cyclic-di-GMP (CDG), is synthesized by bacteria ubiquitously as a secondary messenger (10). CDG is not host cell membrane permeable, rather its accumulation inside of host cells occurs by way of phagocytosis (11, 12). CDG is unique to microbes yet binds numerous mammalian receptors including HCN (13), HpoR (14), Siderocalin (15), and the stimulator of interferon genes (STING) (16). Importantly, CDG has been shown to induce robust type 1 and type 3 interferon (IFN) responses that are critical for anti-viral responses (16-18).

Viral infections are a common cause of asthma exacerbations, though how type 1 and 3 IFNs contribute to asthma pathogenesis is unclear and may be different in specific asthma endotypes. Recently, the global SARS-CoV2 pandemic has led to rapid, broad interest in anti-viral and immunomodulatory strategies to prevent and treat severe disease. Studies in COVID-19 patients have shown that type 1 IFN responses are delayed and/or exaggerated at later phases of severe COVID-19 (19-22). Thus, strategies to control early airway infections through strong local anti-viral responses may be advantageous through modulation of type 1 and 3 IFN and ILC subsets. The effect of cyclic-di-nucleotides on lung ILC responses is unknown. Here, we investigated the role of CDG in fungal allergen and IL-33 driven asthma models.

## RESULTS

### CDG abrogates Alternaria-induced ILC2-driven type 2 airway inflammation and promotes interferon production

To elucidate whether CDG regulates innate lung immunity, we utilized a well-described three-day *Alt*-induced ILC2-driven model of eosinophilic airway inflammation (Figure 1A) (4). Intranasal CDG administration nearly abolished *Alt*-induced bronchoalveolar lavage fluid (BAL) and lung eosinophilia (Figure 1B, C, Supplementary Figure 1A). Concordantly, CDG significantly reduced BAL IL-5 and IL-13 levels (Figure 1D, E). Interestingly, CDG synergistically potentiated *Alt*-induced BAL and lung neutrophil accumulation (Figure 1F, G, Supplementary Figure 1B). In the absence of *Alt*, CDG did not have a significant effect on airway granulocytic infiltration (Figure 1B, C, F, G).

**Figure 1.**
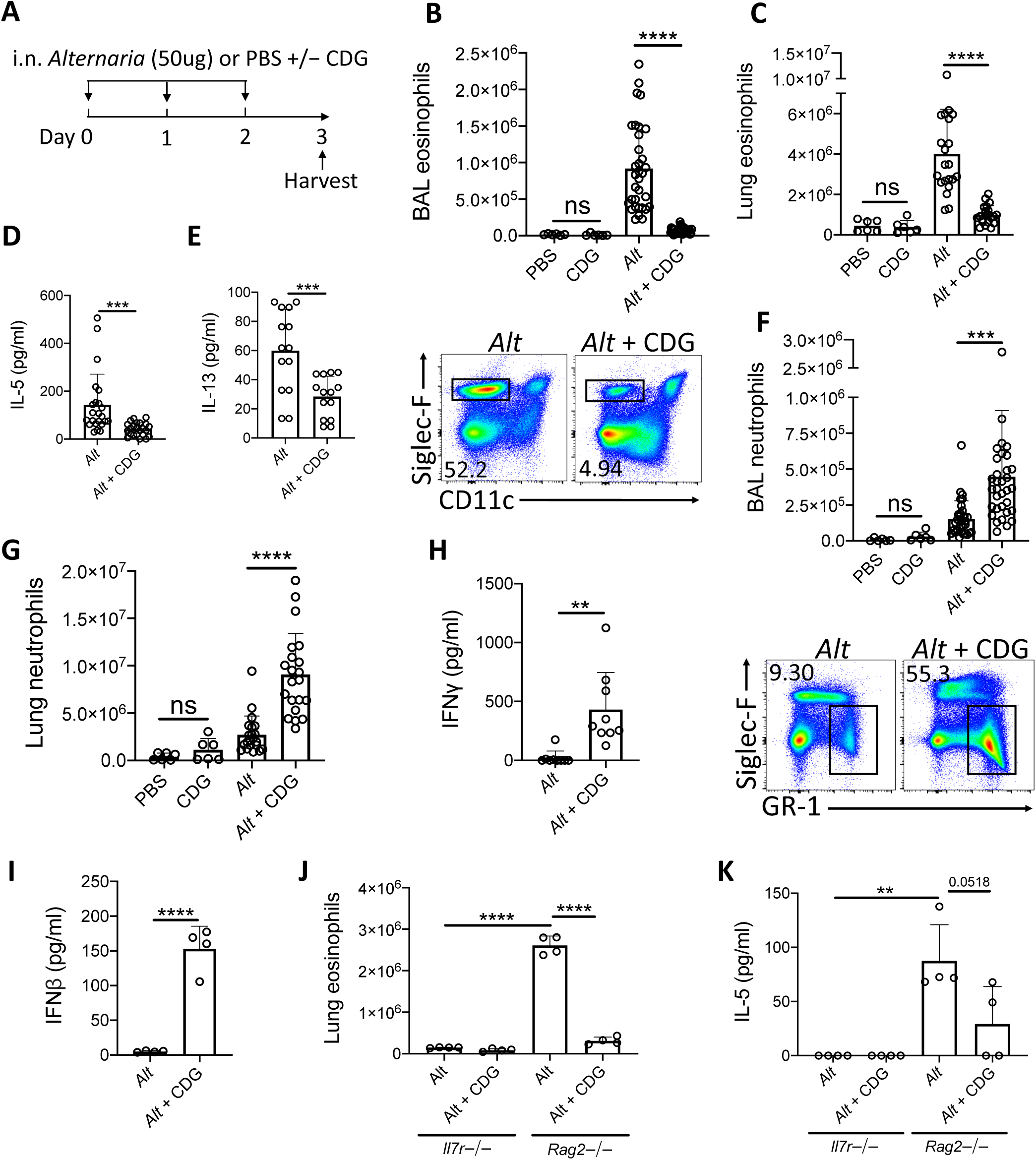
CDG abrogates *Alt*-induced type 2 inflammation and promotes neutrophilia. **(A)** Intranasal challenge mouse model of innate airway inflammation. **(B)** Total BAL eosinophils (top) and representative flow plots (bottom). **(C)** Total lung eosinophils. BAL IL-5 **(D)** and IL-13 **(E)** ELISA. **(F)** Total BAL neutrophils (top) and representative flow plots (bottom). **(G)** Total lug neutrophils. **(H)** BAL IFNγ ELISA. **(I)** BAL IFNβ ELISA 3 hours after first challenge. *Il7r*−/− and *Rag2*−/− lung eosinophils **(J)** and BAL IL-5 ELISA **(K)**. Data shown are representative of 2-10 independent experiments with 2-4 mice per group. *P <.05, **P <.01, ***P <.001, unpaired t test.

Recent studies have shown that ILC2 cytokine production is suppressed by multiple IFNs (23, 24). Therefore, we next asked whether CDG drove IFN production during the *Alt* exposure. Indeed, BAL IFNγ (type 2 IFN) levels increased over 20-fold on average in mice receiving both *Alt* and CDG compared to *Alt* alone (Figure 1H). Further, BAL IFNβ (type 1 IFN) was increased 30-fold three hours following the first intranasal challenge in mice that were CDG challenged (Figure 1I). BAL IFNλ (type 3 IFN) levels were also higher in CDG treated mice at this early timepoint, though the difference was not significant (Supplementary Figure 1C).

To determine whether adaptive or innate lymphocytes, or both, were required for CDG-induced attenuation of type 2 inflammation, we performed experiments with *Rag2*−/− mice that possess ILCs but are deficient in B and T cells as well as *Il7r*−/− mice that lack both ILCs and the majority of T/B cells. Thus, the major difference between these strains is the presence of ILCs (25). No IL-5 and relatively few eosinophils were detectable in the airways of *Il7r*−/− mice (Figure 1J, K, Supplementary Figure 1D). Following *Alt* challenge, lung eosinophil and BAL IL-5 levels were markedly increased in RAG2−/− mice compared to *Il7r*−/− mice. As in wild type mice, CDG nearly abolished *Alt*-induced lung eosinophilia and reduced BAL IL-5 (p=0.0518) in *Rag2*−/− mice. Collectively, these results both demonstrate that CDG combined with *Alt* induces a neutrophilic response associated with increased type 1 and 2 IFNs and suggests that CDG reduces ILC2-driven eosinophilic airway inflammation, independent of adaptive immunity.

### CDG induces a lung ILC2 to ILC1 shift

We next investigated the effect of CDG on ILC2 responses in the three-day *Alt* model. Given the observered increase in BAL IFNγ, we focused on changes in ILC1 responses after CDG exposure. ILC subsets are heterogeneous and show significant plasticity under different conditions (4, 26). Thus, prior to assessing the effect of *in vivo* CDG challenge on ILC subsets, we use transcriptomic analysis to validate the identity ILC1s and ILC2s following *Alt* administration. We reanalyzed our published RNAseq dataset of *Alt* challenged murine lung ST2+CD127+ and ST2−CD127− ILCs (Figure 2A) (4). ST2+CD127+ ILCs highly expressed canonical ILC2 genes (*Klrg1, Il7r, Areg, Il1rl1*, and *Gata3*), whereas ST2−CD127− ILCs highly expressed key ILC1 genes (*Gzma, Gzmb, Irf8, Klrk1*, and *Klrb1*) (Figure 2B). Furthermore, gene ontology pathway analysis indicated that one of the most differentially expressed pathways between ST2+CD127+ and ST2−CD127− ILCs was IFNγ production (Figure 2C). Thus, ST2+CD127+ and ST2−CD127− populations contain strong ILC2 and ILC1 signatures, respectively.

**Figure 2.**
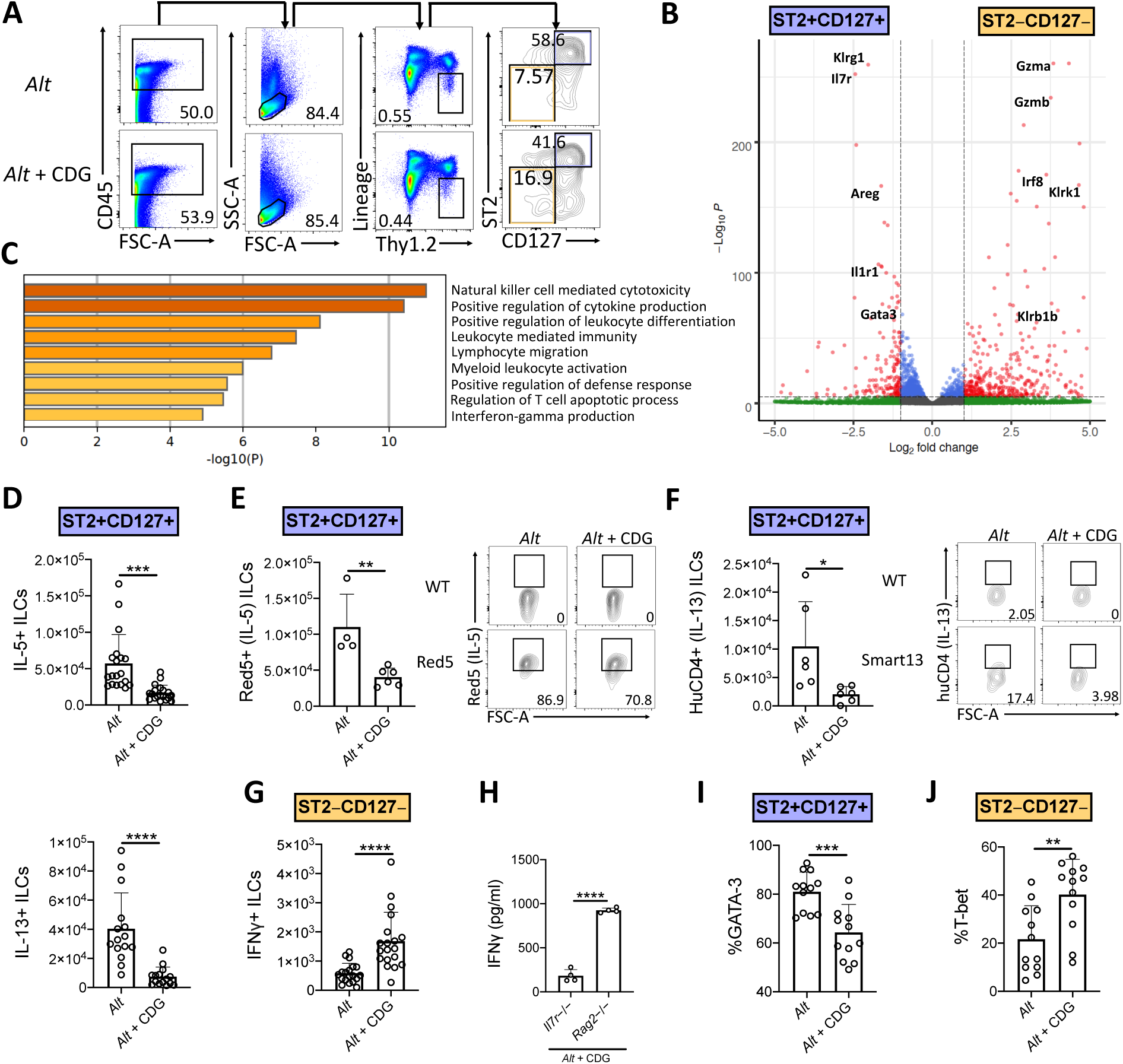
CDG induces ILC2 to ILC1 compartmental changes. Mice were challenged using the same model as in Figure 1A. **(A)** Flow cytometric gating scheme for identification of lung ILC2s and ILC1s in *Alt* and *Alt* plus CDG challenged mice. **(B)** RNAseq volcano plot of differentially expressed genes between *Alt* challenged lung ST2+CD127+ ILCs and ST2−CD127− ILCs. **(C)** Differentially expressed gene ontology pathways in ST2+CD127+ and ST2−CD127− ILC subpopulations. **(D)** Total number of IL-5+ (top) and IL-13+ (bottom) lung ILC2s following PMA/ionomycin stimulation *ex vivo*. **(E)** Total number of Red5+ (IL-5+) lung ILC2s (left) and representative flow plots (right). **(F)** Total number of HuCD4+ (IL-13+) lung ILC2s (left) and representative flow plots (right). **(G)** Total number of IFNγ+ lung ILC1s following PMA/ionomycin stimulation *ex vivo*. **(H)** *Il7r*−/− and *Rag2*−/− BAL IFNγ ELISA. Frequency of GATA-3+ lung ILC2s **(I)** and T-bet+ lung ILC1s **(J)**. Data shown are representative of 2-7 independent experiments with 2-4 mice per group. *P <.05, **P <.01, ***P <.001, unpaired t test.

Intranasal CDG administration drastically reduced the number of *Alt*-induced IL-5+ and IL-13+ lung ILC2s following *ex vivo* PMA/ionomycin stimulation (Figure 2D, Supplementary Figure 2A, B). Because of the relatively artificial nature of *ex vivo* stimulation, we next utilized Red5 IL-5 reporter mice and Smart13 IL-13 reporter mice to provide *in vivo* evidence of CDG-elicited ILC2 suppression (27, 28). Concordantly, we found that CDG attenuated *Alt*-induced lung ILC2 IL-5 and IL-13 production *in vivo* (Figure 2E, F). Contrary to ILC2 responses, the total number of IFNγ+ ILC1s was robustly increased following CDG challenge and *ex vivo* PMA/ionomycin stimulation (Figure 2G, Supplementary Figure 2C). Further, BAL IFNγ levels were markedly increased in *Rag2*−/− mice compared to *Il7r*−/− mice, suggesting that ILC1s likely contribute significantly to CDG-induced IFNγ secretion (Figure 2H).

In light of the CDG-induced shift from an ILC2 to ILC1 response and recent reports of ILC plasticity (26, 29), we further investigated the impact of CDG on ILC identity, activation, and proliferation. CDG decreased lung ILC2 expression of the canonical ILC2 master type 2 cytokine regulator GATA-3 (Figure 2I, Supplementary Figure 2D) and surface markers ICOS and KLRG1 (Supplementary Figure 2E, F). Conversely, CDG increased ILC1 expression of the master type 1 cytokine regulator T-bet by 20% on average (Figure 2J, Supplementary Figure 2G). Further, CDG attenuated ILC2 proliferation, as evidenced by decreased expression of Ki67 (Supplementary Figure 2H), but did not affect ILC2 activation status, as indicated by unchanged CD69 expression levels (Supplementary Figure 2I). However, CDG did not affect ILC1 proliferation (Supplementary Figure 2J) but did significantly increase ILC1 activation status (Supplementary Figure 2K). Taken together, these findings indicate that CDG drives concomitant ILC2 contraction and ILC1 activation.

### CDG induces ILC2 to ILC1 shift during IL-33-driven type 2 lung inflammation

We next asked whether CDG-induced ILC2 inhibition occurred downstream of IL-33 release. To that end, we challenged mice with exogenous IL-33 instead of *Alt*. Consistent with the *Alt* model, CDG severely attenuated IL-33-induced lung eosinophilia (Figure 3A) and robustly increased lung neutrophil accumulation (Figure 3B). Additionally, CDG administration markedly decreased BAL IL-5 (Figure 3C) and increased BAL IFNγ by over 100-fold (Figure 3D). As with *Alt*, CDG decreased the total number of IL-33-induced IL-5+ and IL-13+ lung ILC2s (Figure 3E, F, Supplementary Figure 3A, B) and increased the total number of IFNγ+ ILC1s (Figure 3G, Supplementary Figure 3C). Thus, CDG has effects on IL-33 driven lung responses comparable to those of the fungal allergen *Alt*, suggesting that CDG inhibition of ILC2 responses is not related to an effect on IL-33 levels during *Alt* exposure.

**Figure 3.**
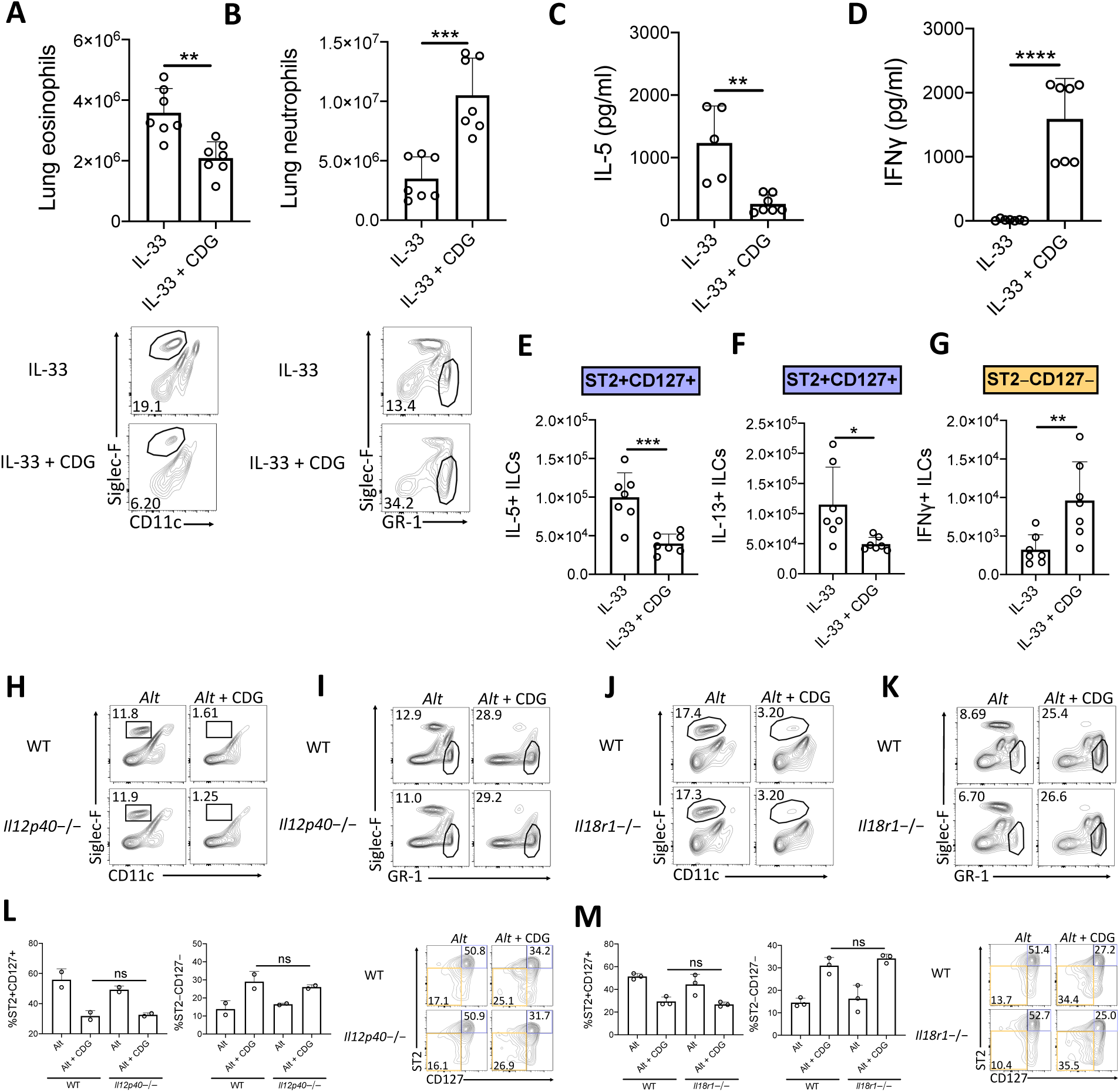
Suppression of type 2 inflammation occurs downstream of IL-33 and is independent of IL-12 and IL-18 signaling. **(A-G)** Mice were challenged using the same model as in Figure 1A but with IL-33 instead of *Alt*. **(A)** Total lung eosinophils (top) and representative flow plots (bottom). **(B)** Total lung neutrophils (top) and representative flow plots (bottom). BAL IL-5 **(C)** and IFNγ **(D)** ELISA. Total number of IL-5+ lung ILC2s **(E)**, IL-13+ lung ILC2s **(F)**, and IFNγ+ lung ILC1s **(G)** following PMA/ionomycin stimulation *ex vivo*. **(H-M)** Mice were challenged using the same model as in Figure 1A. *Il12p40*−/− lung eosinophil **(H)** and neutrophil **(I)** frequencies and *Il18r1*−/− lung eosinophil **(J)** and neutrophil **(K)** frequencies. Data shown are representative of 2 independent experiments with 2-4 mice per group. *P <.05, **P <.01, ***P <.001, unpaired t test.

### IL-12 and IL-18R are dispensable for CDG-induced suppression of type 2 response

The ILC2-driven *Alt* asthma model has been described as largely IL-33 dependent, though conversion of ST2+ ILC2s to ILC1s during lung inflammatory responses to influenza has been shown to be promoted by IL-12 and IL-18 (30, 31). Thus, we investigated whether IL-12 and IL-18 signaling were required for the observed granulocytic shift after addition of CDG to *Alt*. Surprisingly, we found that the levels of airway eosinophilia and neutrophilia, and frequencies of ILC2s and ILC1s, remained constant in mice lacking IL-12 (Figure 3H, I, L, Supplementary Figure 3D, E) and IL-18R (Figure 3J, K, M, Supplementary Figure 3F, G). Collectively, these results suggest that CDG does not regulate ILC2 to ILC1 responses through IL-12 or IL-18.

### CDG suppression of type 2 inflammation and ILC2 responses is STING-dependent

Given the robust airway IFN accumulation induced by CDG, we next investigated whether STING (the <u>st</u>imulator of <u>in</u>terferon <u>g</u>enes) was required for suppression of type 2 lung inflammation and ILC2 responses. Strikingly, through comparison of wild type and STING deficient (*Tmem173*−/−) mice, we found that removal of STING completely rescued CDG-driven abrogation of *Alt*-induced lung eosinophilia and fully abrogated CDG-induced lung neutrophilia (Figure 4A, B). Moreover, lung ILC2 function was completely restored in the absence of STING (Figure 4C), and CDG-driven ILC1 IFNγ induction was entirely STING-dependent (Figure 4D).

**Figure 4.**
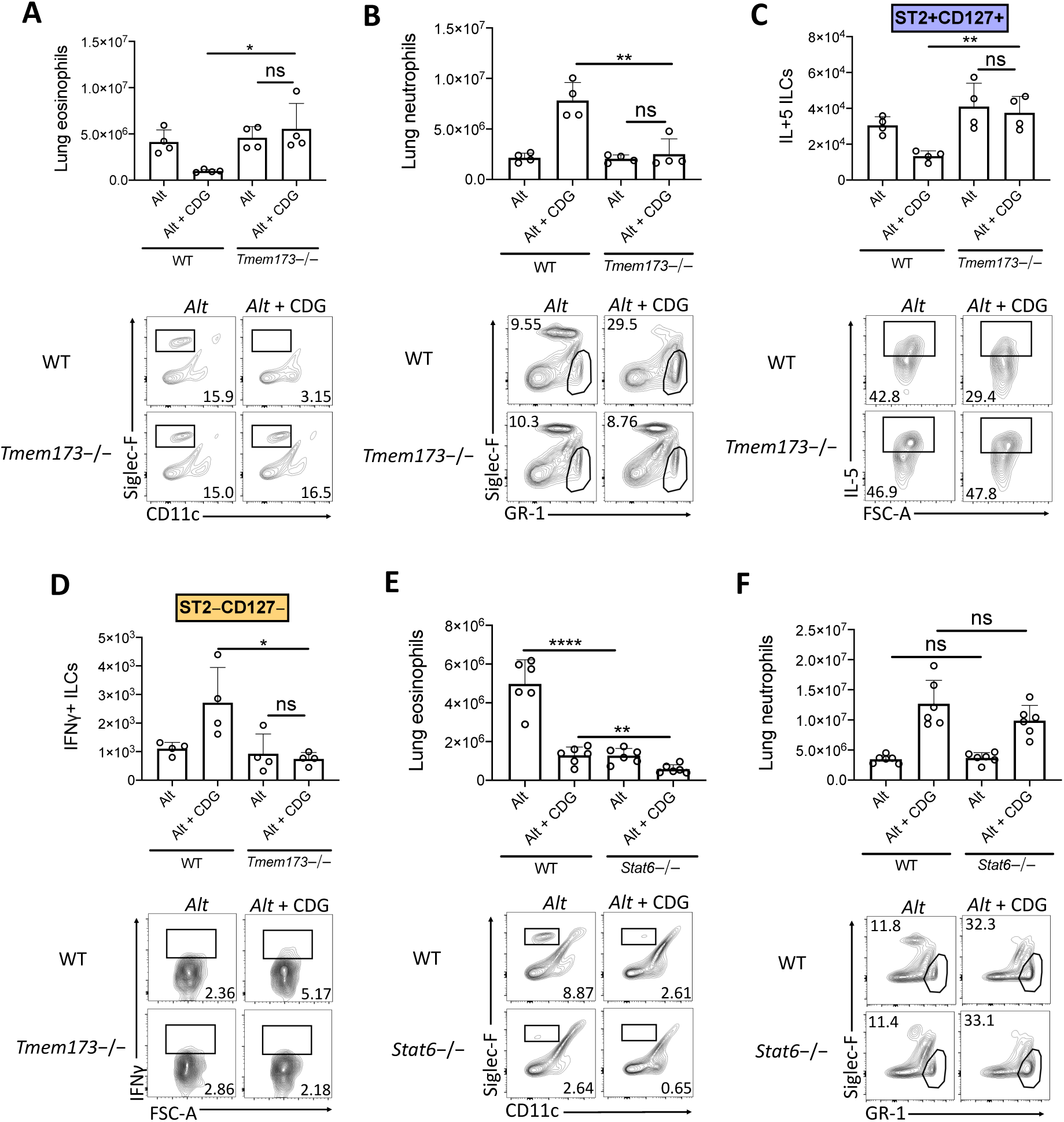
CDG modulation of airway inflammation is STING-dependent and STAT-6 independent. **(A-C)** Wild type and *Tmem173*−/− mice were challenged using the same model as in Figure 1A. **(A)** Total lung eosinophils (top) and representative flow plots (bottom). **(B)** Total lung neutrophils (top) and representative flow plots (bottom). **(C)** Total lung IL-5+ ILC2s (top) and representative flow plots (bottom). **(D-F)** Wild type and *Stat6*−/− mice were challenged using the same model as in Figure 1A. **(D)** Total lung IFNγ+ ILC1s (top) and representative plots (bottom). **(E)** Total lung eosinophils (top) and representative flow plots (bottom). **(F)** Total lung neutrophils (top) and representative flow plots (bottom). Data shown are representative of 2 independent experiments with 2-3 mice per group. *P <.05, **P <.01, ***P <.001, unpaired t test.

### STAT6 is not required for neutrophilic airway inflammation induced by CDG

A recent study demonstrated that STAT6 was required for STING-dependent chemokine production (32). Therefore, we sought to determine whether STAT6 was required for neutrophil accumulation in lung tissue following CDG challenge. Consistent with previous reports, *Alt*-induced eosinophilia was highly dependent upon STAT6 (Figure 4E) (25). Notably, we found that residual lung eosinophil levels following CDG administration were further reduced in the absence of STAT6. Unlike eosinophils, neutrophils accumulated in the airway independent of STAT6 (Figure 4F). These results demonstrate that while STAT6 regulates innate type 2 lung inflammation, likely through chemokine expression, it has no role in the STING-induced effects on neutrophil accumulation (25, 33).

### Type 1 IFN signaling is partially required for CDG induced immunomodulation

Type 1 IFNs are thought to be the major products of <u>st</u>imulator of <u>in</u>terferon <u>g</u>enes (STING) activation. To confirm that CDG-induced type 1 IFN production was responsible for the observed immunomodulation, we compared type 1 IFN signaling deficient (*Ifnar1*−/−), wild type, and *Tmem173*−/− mice. Surprisingly, *Ifnar1*−/− mice did not phenocopy *Tmem173*−/− mice. Unlike STING deficiency, removal of type 1 IFN signaling only partially rescued CDG-induced abrogation of BAL eosinophilia (Figure 5A) and did not fully prevent CDG-induced BAL neutrophilia (Figure 5B). Moreover, as in wild type mice, the frequency of ILC1s was significantly increased following CDG challenge in mice deficient in type 1 IFN signaling; whereas, the frequency of ILC2s remained constant in *Ifnar1*−/− mice, as in those lacking STING (Figure 5C). Thus, type 1 IFNs are only partially required for CDG-induced immunomodulation, indicating the observed shifts in peribronchial inflammation and ILCs depend on an alternative mediator downstream of STING.

**Figure 5.**
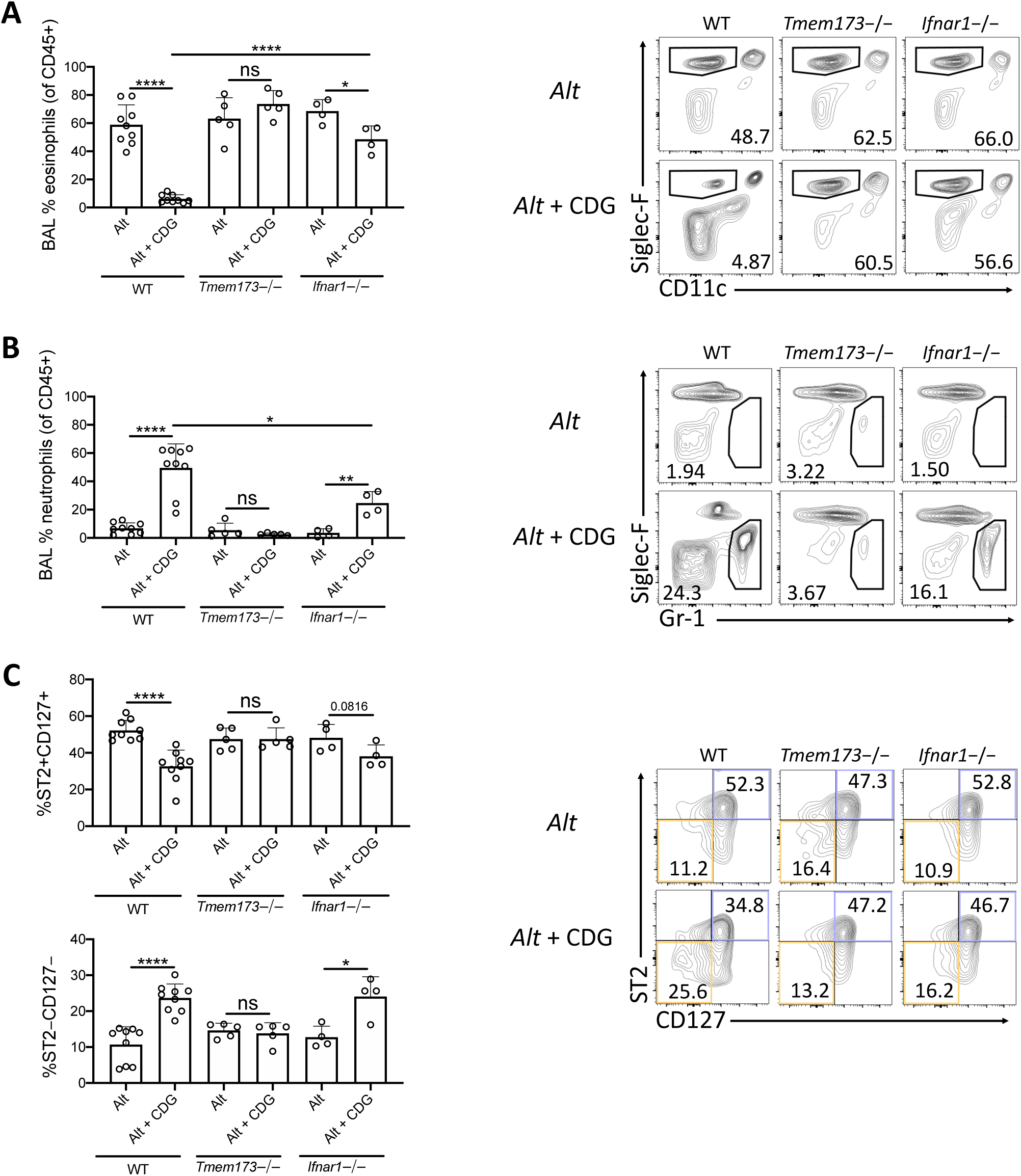
Type 1 IFN signaling is partially required for CDG induced immunomodulation. Wild type, *Tmem173*−/−, and *Ifnar1*−/− mice were challenged using the same model as in Figure 1A. **(A)** Frequency of BAL eosinophils (left) and representative plots (right). **(B)** Frequency of BAL neutrophils (left) and representative flow plots (right). **(C)** Frequency of lung ST2+CD127+ and ST2−CD127− ILC subpopulations (left) and representative flow plots (right). Data shown are representative of 5 independent experiments with 1-3 mice per group. *P <.05, **P <.01, ***P <.001, unpaired t test.

## DISCUSSION

Asthma is largely a type 2 inflammatory airway disease and is associated with bacterial and viral infections, mitochondrial stress, and host cell death, which all lead to the accumulation of the danger-associated cyclic-di-nucleotides (5-8, 34). ILC2s promote type 2 inflammation in experimental asthma models and likely contribute to airway inflammation and hyperresponsiveness in humans (35). Here, we investigated how *in vivo* administration of the cyclic-di-nucleotide CDG regulates ILC2s and innate type 2 inflammation. Our novel findings show that intranasal CDG challenge nearly abolished *Alt*-induced airway eosinophilia but promoted neutrophil accumulation. Congruently, CDG drove contraction of IL-5+ and IL-13+ lung ILC2s and expansion of IFNγ+ ILC1s. Mechanistically, we found that CDG modulated ILC responses downstream of *Alt*-induced IL-33 release and that the effect of CDG on airway inflammation and ILC function was entirely STING-dependent and partially type 1 IFN-dependent. STING’s involvement in pathogen responses, as well as activation by cytosolic mitochondrial DNA (from damaged mitochondria) and autophagy, could represent a critical, common pathway in the development of mixed airway inflammatory responses often found in asthma (8, 36-40).

While type 1 IFNs are the major products of STING activation, STING agonism has also been shown to induce the production of several inflammatory mediators including type 3 IFN, IL-1β, IL-6, TNFα, CCL2, and CCL20 (41-44). In addition to elevated levels of type 1 IFNs, we observed an increase in type 3 IFN following CDG administration, though it failed to reach significance. Type 1, 2 and 3 IFNs are known to inhibit ILC2 function and restrict type lung 2 inflammation (23, 24, 45, 46), however the effect on development of ILC1 accumulation by IFNs is not well studied. Taken together with recent reports, our findings suggest a model in which CDG drives STING-dependent early production of type 1 and 3 IFNs, which suppress ILC2s and promote ILC1 IFNγ that further suppresses ILC2s. Because the effect of CDG on *Alt*-induced inflammation we observed was only partially dependent on type 1 IFN signaling, non-IFN mediators downstream of STING likely contribute to CDG-induced immunomodulation.

The effect of CDG on airway eosinophil and neutrophil levels we report here agrees with a previous report in which CDG was administered in an adaptive Th2 cell-driven chronic asthma model, though ILC changes were not assessed (47). Further, the ILC compartmental changes we observed mirror influenza virus-induced ILC2 to ILC1 plasticity described in a recent study (30). However, that study demonstrated that IL-12 and IL-18 drove the attenuation of type 2 cytokine production, whereas we found that CDG abrogated type 2 inflammation independent of IL-12 and IL-18 signaling and dependent on STING and type 1 IFNs. Collective, these findings indicate that there are multiple ways to induce an ILC2 to ILC1 shift in the lungs.

Interestingly, a recent investigation into the role of STING in the gastrointestinal mucosa revealed a nearly two-fold reduction in ILC2 frequency and IL-4 and IL-13 levels, and a three-fold increase in ILC1 frequency, in STING deficient mice (48). Conversely, we found markedly increased airway ILC2s and type 2 cytokines, and decreased ILC1s and type 1 cytokines, in STING deficient mice following *Alt* and CDG challenge. Such highly divergent findings suggest the presence distinct, tissue-specific mechanisms by which STING controls ILC responses. These results are consistent with recent work demonstrating tissue specific ILC responses (49).

The findings we report here might be leveraged for developing treatments to induce early, local control of respiratory viruses, including SARS-CoV-2, in which type 1 and 3 IFN control of virus may be critical for prevention of late complications in severe disease (19, 21). RNA viruses, such as coronaviruses, induce STING downstream of RIG-I and MAVs signaling (50). Importantly, however, coronaviruses have evolved mechanisms for inhibiting STING activation. For example, coronaviruses encode proteases that antagonize STING and prevent downstream IFN production (51, 52). Interestingly, bats, which harbor coronaviruses yet show no signs of immunopathology, constitutively express IFN and were recently shown to possess a distinct STING from that of other mammals (53). When bat STING protein was humanized with a single amino acid substitution, viral tolerance was lost, suggesting a critical role for STING activation and IFN expression in controlling viral replication. Additionally, patients with severe COVID-19 have impaired type 1 IFN responses (21), and early administration of the type 1 interferon IFN-α2b led to reduced in-hospital mortality in COVID-19 patients (22). Collectively, our work, coupled with these reports, suggests that CDG, or other human STING agonists, may be a potential low-cost, local respiratory therapeutic for early treatment of COVID-19 following SARS-CoV2 infection through broad generation of anti-viral type 1 and 3 IFNs.

Our study is limited in that mechanisms downstream of STING that account for airway neutrophil and ILC1 accumulation remain unclear. Consistent with previous reports, we showed that lung eosinophilia is dependent on STAT6 (25). A prior study showed that STAT6 was required for STING-mediated chemokine production and immune cell recruitment (32); however, in our study, CDG-induced airway neutrophil accumulation occurred independent of STAT6, which suggests the existence of an alternative pathway by which immune cells accumulate in the presence of STING agonism. Future studies investigating this distinct mechanism are warranted.

In summary, our work demonstrates that CDG drives STING-dependent IFN production, ILC1 activation and accumulation, as well as ILC2 suppression and abrogation of innate type 2 innate airway inflammation. The knowledge set forth in this report is critical for understanding how allergic airway disease pathogenesis may be impacted by lung insults due to cellular stress, bacterial infection, or viral infections such as SARS-CoV2.

## MATERIALS AND METHODS

### Mice

6- to 12-week-old female C57BL/6 WT mice were obtained from The Jackson Laboratory (Bar Harbor, Me). *Rag2*−/−, *Il7r*−/−, *Il12p40*−/−, *Ifnar1*−/−, *Tmem173*−/−, and *Stat6*−/− mice were obtained from The Jackson Laboratory and bred in-house. Red5 mice were obtained from Dr. Nunzio Bottini (UCSD), originated from Dr. Richard Locksley (UCSF), and bred in-house (28). Smart13 mice were obtained from The Jackson Laboratory, originated from Dr. Richard Locksley (UCSF), and bred in-house (27). *Il18r1*−/− mice were obtained from Dr. Hal Hoffman (UCSD) and bred in-house. All animal experiments were approved by the University of California, San Diego Institutional Animal Care and Use Committee.

### In vivo Alternaria models

WT, gene knockout, and cytokine reporter mice were challenged intranasally with PBS, 5ug CDG, 50ug of *Alt* extract (Greer, lot number 299382), or 5ug CDG and 50ug *Alt* extract in 40uL every 24 hours for 3 days and euthanized 24 hours following the third challenge for BAL collection and lung tissue harvest. A subset of experiments was performed with only the latter two experimental groups. Further, in select experiments, *Alt* was replaced with exogenous IL-33 (500ng). For experiments investigating early cytokine release, mice were euthanized 3 hours following the first challenge. BAL was performed with 2% BSA in PBS; the first draw was 500uL and draws 2-5 were 600uL. BAL was centrifuged at 1500 rpm for 5 minutes at 4C and the supernatant was stored at −20C for ELISA and cells were counted and phenotyped using flow cytometry. Lungs were digested using the Mouse Lung Dissociation Kit (Miltenyi Biotec) according to the manufacturer’s protocol, filtered with a 40um mesh, and cells were counted and phenotyped using flow cytometry.

### Flow cytometry

Mouse BAL and lung cells were suspended in a solution of 2% FBS and 0.01% sodium azide in PBS and counted on a Novocyte (Acea Biosciences). Cells were incubated with an unconjugated mAb to CD16/CD32 for 10 minutes at 4C to block non-specific Fc receptor binding and then incubated for 30 minutes with fluorochrome conjugated antibodies at 4C. All antibodies were purchased from BioLegend unless otherwise noted. To identify eosinophils (CD45+Siglec-F+CD11c−) and neutrophils (CD45+GR-1+Siglec-F−), BAL or lung cells, or both, were stained with PerCP-conjugated anti-CD45.2, PE-conjugated anti-Siglec-F (BD), FITC-conjugated anti-CD11c, and APC-conjugated anti-GR-1. For the identification of ILCs (CD45.2+lineage−Thy1.2+ lymphocytes), ILC2s (CD45.2+lineage− Thy1.2+ST2+CD127+), and ILC1s (CD45.2+lineage-Thy1.2+ST2−CD127−) lung cells were incubated with FITC–conjugated lineage cocktail (anti-CD3e, anti-GR-1, anti-CD11b, anti-B220, anti-Ter119), anti-CD11c, anti-NK1.1, anti-CD5, anti-FcεR1, anti-TCRβ, and anti-TCRγδ; PerCP-conjugated anti-CD45.2; eFluor 450-conjugated anti-Thy1.2 (ThermoFisher); APC-conjugated anti-ST2; and PE-Cy7-conjugated anti-CD127. In select experiments, cells were also stained with PE-conjugated anti-CD69, anti-ICOS, or anti-KLRG1. For transcription factor staining, surface-stained cells were fixed and permeabilized with a transcription factor staining kit (ThermoFisher) according to the manufacturer’s protocol and stained for 30 minutes at 4C with PE-conjugated anti-GATA-3 (ThermoFisher), anti-Ki67 (ThermoFisher), or anti-T-Bet.

For ILC cytokine staining, lung cells from WT and knockout mice were incubated for 3 hours at 37C with a phorbol 12-myristate 13-acetate, ionomycin, brefeldin A, and monensin cell stimulation cocktail (ThermoFisher) in RPMI 1640 media supplemented with penicillin/streptomycin, 10% FBS, glutamine, and 2-mercaptoethanol (ThermoFisher). Stimulated cells were surfaced stained for ILC2s and ILC1s as described above, fixed and permeabilized with an intracellular cytokine staining kit (BD) according to the manufacturer’s protocol, and stained for 30 minutes at 4C with PE-conjugated anti-IL-5 or anti-IL-13 and APC-Cy7-conjugated anti-IFNγ.

Reporter mice were used to assess cytokine staining directly *ex vivo* without stimulation. To visualize *in vivo* IL-13 in Smart13 mice, human CD4 was stained with PE-conjugated anti-human CD4.(27) To visualize *in vivo* IL-5 in Red5 mice, the PE channel was used without any requisite staining.(28) Finally, samples were analyzed with a Novocyte (Acea Biosciences) flow cytometer.

### ELISA

ELISAs for IL-5 (R&D), IL-13 (R&D), IFNγ (ThermoFisher), IFNβ (R&D), and IFNλ (R&D) were performed on BAL supernatants according to the manufacturers’ protocols. ELISA plates were read on a model 680 microplate reader (Bio-Rad) at 450 nm.

### RNAseq

Publicly available RNA sequencing of mouse lung ILC2s and ILC1s (GEO: GSE136156) was downloaded. Reads were then aligned to reference genome mm10 using TopHat. DUST scores were calculated with PRINSEQ Lite and low complexity reads (DUST > 4) were removed from the BAM files. Read counts to each genomic feature are obtained with the HTSeq count program. Differentially expressed genes were identified with DESeq2 and pathway analysis was performed with Metascape.

### Data analysis and statistics

Flow cytometry data were analyzed using FlowJo version 10.4.1 (Flowjo). For all experiments (except RNAseq), statistical analysis was performed using GraphPad Prism software (GraphPad) using unpaired t-tests (2-tailed). P values of less than.05 were considered statistically significant.

## ABBREVIATIONS

(ILC): Innate lymphoid cell
(ILC2): group 2 innate lymphoid cell
(ILC1): group 1 innate lymphoid cell
(*Alternaria alternata*): *Alt*
(CDN): cyclic-di-nucleotide
(CDG): cyclic-di-GMP
(STING): stimulator of interferon genes
(IFN): interferon
(BAL): bronchoalveolar lavage

## AUTHOR CONTRIBUTIONS

K.J.C, J.H.B., M.B.A., L.H.N., L.D.L, A.S.L, A.N.S. performed the experiments, K.J.C. and T.A.D. designed and interpreted the experiments, and K.J.C. and T.A.D. wrote the manuscript.

## ACKNOWLEDGEMENTS

We thank Dr. Hal Hoffman and Dr. Nunzio Bottini for the *Il18r1*−/− and Red5 mice, respectively.

## DATA AVAILABILITY STATEMENTS

No data related to this manuscript were deposited in a public database.

## ETHICS STATEMENT

The animal research reported here was conducted following approval by the University of California, San Diego Institutional Animal Care and Use Committee.

## FIGURE LEGENDS

**Supplementary Figure 1.**
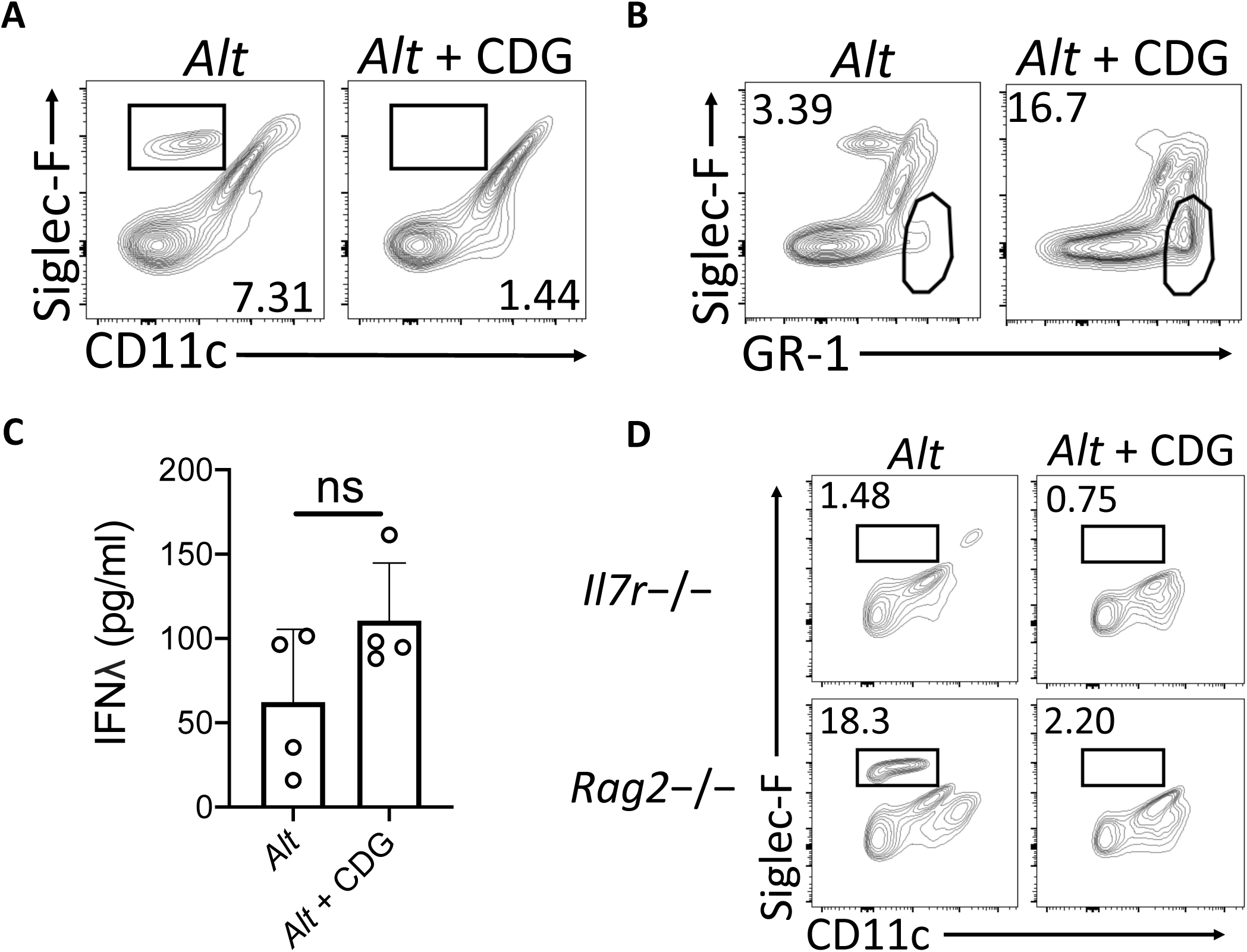
CDG abrogates *Alt*-induced type 2 inflammation and increases neutrophilia. Mice were challenged using the same model as in Figure 1A. Representative flow plots for eosinophil **(A)** and neutrophil **(B)** identification in wild type mice. **(C)** BAL IFNλ ELISA. **(D)** Representative flow plots for eosinophil identification in *Rag2*−/− and *Il7r*−/− mice. Data shown are representative of 2-10 independent experiments with 2-4 mice per group. *P <.05, **P <.01, ***P <.001, unpaired t test.

**Supplementary Figure 2.**
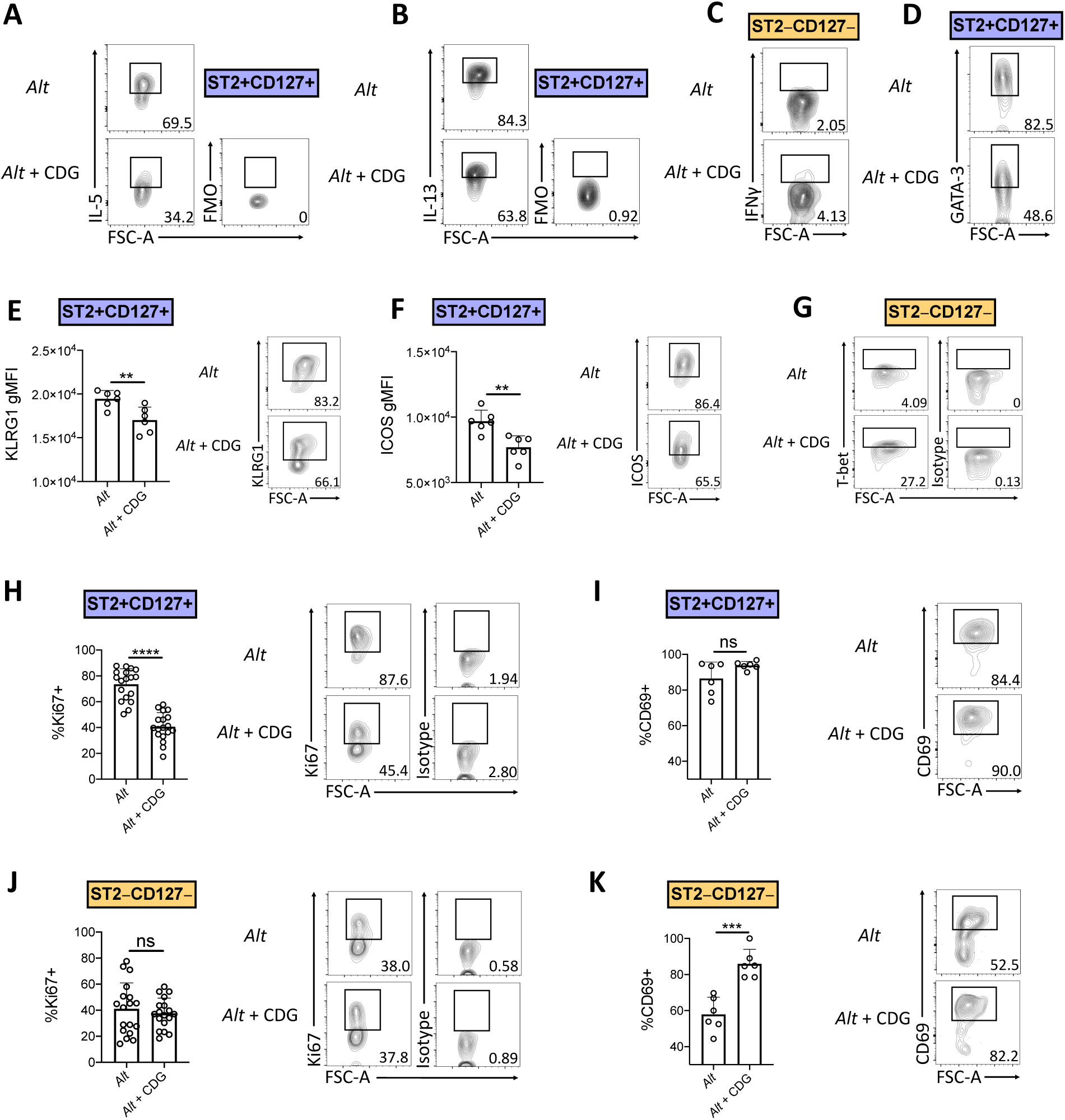
CDG induces innate lymphoid cell compartmental changes. Mice were challenged using the same model as in Figure 1A. Representative flow plots for lung ILC2 IL-5 **(A)**, ILC2 IL-13 **(B)**, ILC1 IFNγ **(C)** and ILC2 GATA-3 **(D). (E)** Geometric mean fluorescent intensity (gMFI) of lung ILC2 KLRG1 (left) and representative flow plots (right). **(F)** gMFI of lung ILC2 ICOS (left) and representative flow plots (right). **(G)** Representative flow plots for lung ILC1 T-bet. **(H)** Frequency of Ki67+ lung ILC2s (left) and representative flow plots (right). **(I)** Frequency of CD69+ lung ILC2 (left) and representative flow plots (right). **(J)** Frequency of Ki67+ lung ILC1 (left) and representative flow plots (right). **(K)** Frequency of CD69+ lung ILC1 (left) and representative flow plots (right). Data shown are representative of 2-7 independent experiments with 2-4 mice per group. *P <.05, **P <.01, ***P <.001, unpaired t test.

**Supplementary Figure 3.**
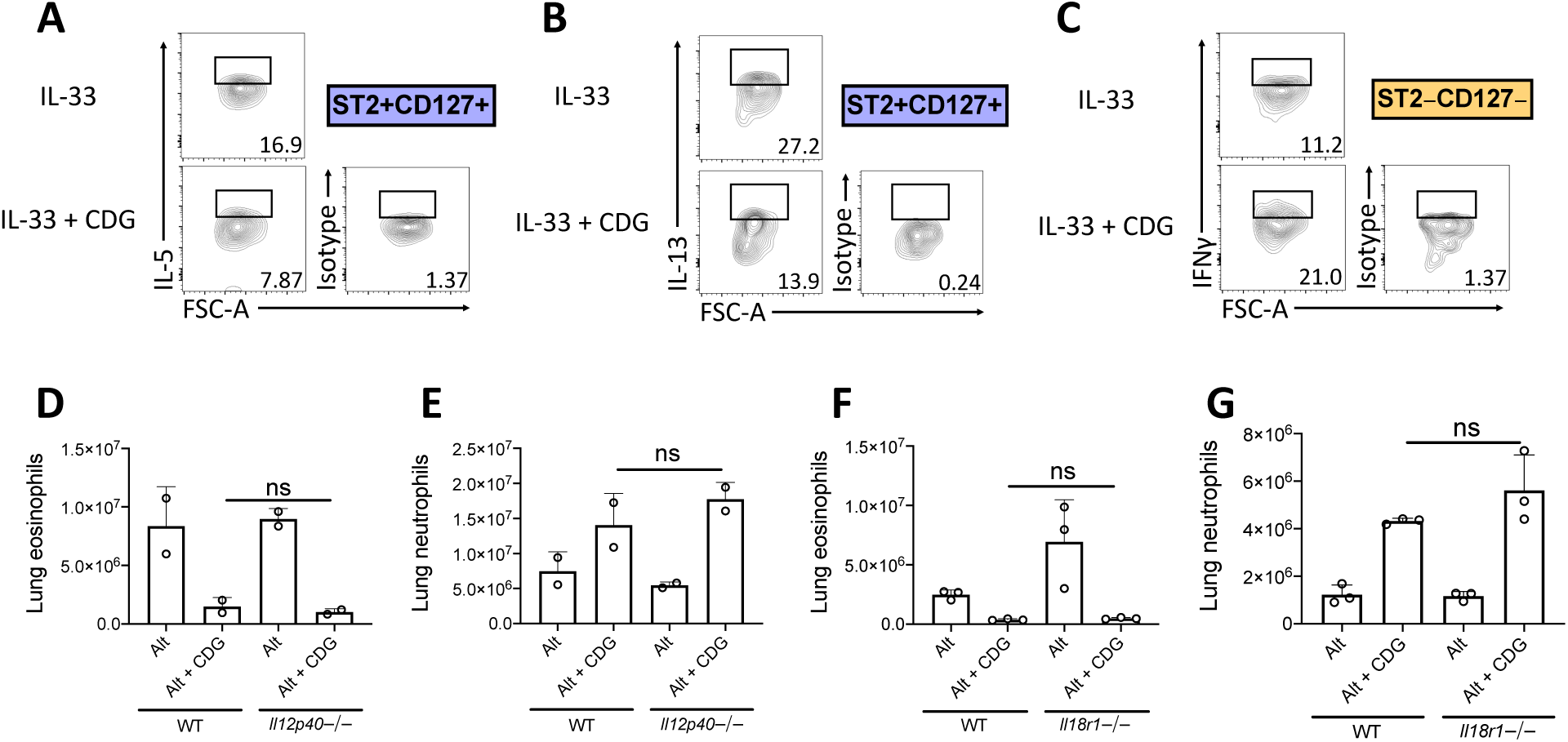
Suppression of type 2 inflammation occurs downstream of IL-33 and independent of IL-12 and IL-18 signaling. **(A-C)** Mice were challenged using the same model as in Figure 1A but with IL-33 instead of *Alt*. Representative flow plots for lung ILC2 IL-5 **(A)**, ILC2 IL-13 **(B)**, and ILC1 IFNγ **(C). (D-G)** Mice were challenged using the same model as in Figure 1A. Total number of lung eosinophils **(D)** and neutrophils **(E)** in wild type and *Il12p40*−/− mice. Total number of lung eosinophils **(F)** and neutrophils **(G)** in wild type and *Il18r1*−/− mice. Data shown are representative of 2 independent experiments with 2-4 mice per group. *P <.05, **P <.01, ***P <.001, unpaired t test.

